# Experimentally infection of Cattle with wild types of Peste-des-petits-ruminants Virus – Role in its maintenance and spread

**DOI:** 10.1101/331827

**Authors:** Emmanuel Couacy-Hymann, Valère K. Kouadio, Mathurin Y. Koffi, Arsène Mossoum, Antoinette Kouassi, Krou Assemian, Privat H. Godji

## Abstract

PPR is a common and dreadful disease of sheep and goats in tropical regions caused by PPRV which can infect also cattle without any clinical signs but show a seroconversion. However the epidemiological role of cattle in the maintenance and spread of the disease is not known. For the purpose of the present study, cattle were infected with a wild candidate from each of the four lineages of PPRV and placed in separate boxes. Then naive goats were introduced in each specific box for the 30 days duration of the experiment. The results showed that no clinical signs of PPR were recorded from these infected cattle along with the in-contact goats. The nasal and oral swabs remainend negative. However, animals infected with wild types of PPRV from lineages 1, 3, 4 seroconverted with high percentage inhibition (PI %= values. Only two animals out of three with the Nigeria 75/3 strain of lineage 2 (mild strain) did elicit a production of specific anti-PPR antibodies in those cattle but with PI% values around the threshold of the test. Our findings confirm that cattle are dead end hosts for PPRV and do not play an epidemiological role in the maintenance and spread of PPRV. In a PPR surveillance programme, cattle can serve as indicators of PPRV infection.

**Importance:** Peste-ds-ptetis-ruminants (PPR) is a major Transboundary Animal disease (TADs) in the tropical regions which is spreading extensively nowadays to southern and northern of Africa, Turkey in Europ and southwest Asia. PPR virus is very close related to Rinderpest virus (RPV) which has been eradicated from the world. Today FAO, WOAH / OIE and the scientific community have elected PPR to be the second animal disease to be eradicated through The PPR Global Eradication Programme (GEP-PPR). Since PPR infects cattle without any clinical signs but they seroconvert, it is important to explore the role of cattle in the maintenance and spread of PPRV to better understand the epidemiology of the disease which wll help in the the GEP-PPR.

## Introduction

Peste Des Petits Ruminants (PPR) is a serious and contagious plague of small ruminants, mostly sheep and goats, in many developing countries in Africa, near and Middle-East and southern Asia (1, 2)). Within Africa, PPR has now extended to southwards in Tanzania, Democratic Republic of Congo and Angola (3, 4). Outbreaks of PPR have been also reported across North Africa including Algeria, Morocco, Tunisia (5, 6) along with the European part of Turkey (7). In southwest Asia, China has reported PPR spread all over the country starting during year 2007 in Tibet region (8). The current spread of PPR over large geographical areas is certainly a result of intensified animal movement and trade but may also be due to the eradication of RPV that affected small ruminants and induced immunity against PPR. Animal of all ages are susceptible and the transmission route remains oral and respiratory secretions following close contact between infected and naive population (9).

The causative agent, Peste Des Petits Ruminants Virus (PPRV) is a negative-stranded RNA virus with a monosegmented genome of length 15,948 and containing six genes encoding six structural proteins. It belongs to family *Paramyxoviridae* and the genus *Morbillivirus* together with Rinderpest Virus (RPV), Measles Virus (MV), Canine Distemper Virus (CDV) and marine mammalian Morbilliviruses (10, 11). There are four lineages of PPRV based on the differentiation determined by the sequence comparison of a small region of the F gene (12) or the N gene (13). However, it has been demonstrated recently that the N gene is more divergent therefore more suitable for phylogenetic distinction between closely related PPRV viruses (14).

The disease is highly contagious and case fatality rates in some outbreaks can approach 90% in susceptible populations and, as a consequence of the effects of epidemics, the local and rural economies of the affected countries can be devastating (15, 16). Nowadays there are efficient attenuated vaccines to be used to prevent this disease and to control its extension (17, 18).

PPRV infects also cattle but only causes disease in small ruminant species while a specific seroconversion to PPR is observed in cattle (19). However, a high mortality of domestic buffaloes (*Bubalus bubalis*) was noted in India caused by an infection with PPRV (20). Even though this situation has not been reported again, there is a necessity to clarify it experimentally and by collection of data from rural communities where mixed species (cattle and small ruminants) graze together.

The present study aimed to investigate the epidemiological role of cattle in the maintenance and spread of PPRV among cattle and small ruminants‘populations.

## Material and Methods

### Animal

Cattle: 15 individuals (N’dama breed), two-three years old, were randomly selected from a farm belonging to the Centre for Research in Agronomy (CNRA – La Mé), located at approximatively, 30 kms from Abidjan. They were tested as being negative for antibodies to PPRV using a competitve ELISA (21). Then they were housed in boxes with separate feeding and drinking tanks. These animals were treated with the anthelmintic Albendazole (10mg/kg) two times during the acclimatisation period lasting ten days.

Goats: 15 West African dwarf goats, randomly selected from the same centre (including seven control goats), aged one - two years, which were tested negative for the presence of antibodies against PPR by PPR competitive ELISA (c-ELISA) (21), were used for the study. Each animal was treated with the anthelmintic Albendazole (7.5 mg/kg) two times during the acclimatisation period (including infected control and uninfected control goats).

After 10 days for the acclimatisation period, the 15 individuals cattle were, at random, divided in four groups of three each with the fifth group (conrol) having also three animals. Each group was randomly assigned to one specific box corresponding to a specific PPRV lineage (Table1).

### All animals in the experiment were earmarked with a unique identification number

#### Virulent isolates used in challenge

Four virus isolates were obtained from the virus bank of CIRAD-Montpellier (France) representing viruses from different geographical regions and belonging to different lineages based on the sequences of their nucleoprotein (NP) gene (14, 22): CIV89 (Lineage 1), Nigeria 75/3 (lineage 2), Ethiopia (lineage 3), India-Calcutta (lineage 4).

### Virulent challenge

Each individual cattle (except uninfected controls) was infected subcutaneously with 1 mL of the various challenge viral suspensions, at a concentration of 10^3^ TCID_50_/mL. Animals were kept separately in boxes. Three cattle were not infected and used as controls.

Infected Control goats: two goats were infected subcutaneously with 1 mL of CIV89 strain, at a concentration of 10^3^ TCID_50_/mL and two goats infected with India-Calcutta strain at the same concentration.

### Uninfected control goats: three goats were not infected

Twenty four hours (24h) after the virulent challenge of cattle, randomly two uninfected and naive goats were introduced into each box already containing infected cattle with a sepecific challenge strain of PPRV.

Infected goats with CIV89 and with India-Calcutta strains respectively, were kept in separated boxes in another animal building. Uninfected control goats were kept in a different box in the same building.

An attendant was assigned to each box to feed and water the infected and control animals. Animals were examined daily for classical signs of PPR and body temperatures were recorded for first ten days post infection (pi) then only for clinical examination up to 30 days pi for cattle.

The study was approved by the Ethics Committee of LANADA – Abidjan and by the National Ethics Committee - Ivory-Coast. In addition the principal investigator and corresponding author was certified from the **International Council for Laboratory Animal Science** (ICLAS).

### Sample collection

Serial bleeding was performed on all animals at: day0, day2, day5, day7, day9 then day15, day30 post infection (end of the study) for cattle and in-contact goats and up to day8 for infected control goats. Serum was separated and samples stored at −20°C until examined.

Swab: ocular and nasal swabs were collected at day0, day2, day5, day7, day9 then day15, day21 and day30 post infection, for cattle and in-contact goats. Individual sterile swabs were used in the present experiment. In the Centre, collected swab samples were kept in liquid nitrogen to prevent any degradation of biological materials. At the laboratory, swabs were transferred to a −80°C freezer until used for analysis.

### Serological test

A competitive ELISA (cELISA) kit (CIRAD-Montpellier, France), based on a recombinant NP was used to detect specific antibodies against PPR (21)) following recommended protocols. Fifty microlitres were used throughout. Maxisorp 96-wells plates were coated with the recombinant NP antigen diluted 1/1600 in PBS (0.01 M, pH 7.2–7.4) and incubated at 37°C for 1 h on an orbital shaker. After a cycle of three washes in phosphate buffered saline (PBS; 1/5, 0.05% Tween 20), test serum (5 µL), was added to 45µL of blocking buffer (PBS 0.01 M. pH 7.2–7.4; 0.05% Tween 20 (v/v); 0.5% negative sheep serum (v/v)) followed immediately by the addition of 50µL of the specific monoclonal antibody (Mab) against the PPRV NP at a dilution of 1/100 in blocking buffer. Control sera included were,strong positive, weak positive, negative and a Mab control (0% competition). The plates were incubated and washed as above. Anti-mouse horse radish peroxidase enzyme conjugate (DAKO A/S), diluted 1/1000 in blocking buffer, was added and plates incubated as before. The plates were washed and 50µL of substrate/ chromogen (H2O2/OPD) were added and the colour allowed to develop for 10 min, after which time any reaction was stopped by the addition of 50µL of sulphuric acid (1 M.). Plates were read on an ELISA reader (Multiskan MK II) at an absorbance of 492 nm. Optical density (OD) readings were converted to percentage inhibition (PI) values using the following formula:

PI% = 100 (OD in test well / OD in 0% control well) x100.
PI% values greater than or equal to 50% were considered positive

### Single stranded cDNA synthesis and PCR technique

Oral and nasal swabs were processed as described (13). The procedure for RNA isolation was as recommended by the manufacturer, using the RNeasy Mini Kit (Qiagen, Germany). The RNA was eluted in 50 μL of nuclease-free water. The RT step was performed by using random hexamer primers (Introgen, Carlsbad, CA., USA) with 10 μL of extracted RNA and the First-strand cDNA Synthesis Kit (GE Healthcare Europe GmBH, Orsay,France) as recommended by the manufacturer’s protocol. Then, 5 μL of the cDNA obtained was used as the template for the PCR step in a 200 µL thin wall tube. The PCR was carried out using the Gene Amp PCR system 2400(Perkin-Elmer, Applied Biosystems, Paris, France) using a 50 µL reaction mixture with the specific set of primers NP3 (forward:5’ – TCT CGG AAA TCG CCT CAC AGA CTG) and NP4 (reverse: 5’ – CCT CCT CCT GGT CCT CCA GAA TCT) as previously outlined (13) targeting a fragment of 350 bp on the nucleoprotein (NP) with the following programme: an initial denaturation step at 95°C for 5 min followed by five cycles with denaturation at 94°C for 30 sec, annealing at 60°C for 30 sec and the extension at 72°C for 30 sec. Then the amplification process continued for 30 cycles more but in which the annealing temperature was reduced to 55°C. The amplification reaction was terminated by a final extension of 10 min at 72°C. Negative and positive controls were included in all experiments.

## Results

Clinical response of goats (Infected Control Goats) to infection with PPRV CIV 89 and India-Calcutta isolates

For both PPRV strains used, the infected goats developed pyrexia after an incubation period of 2–7 days, with rectal temperatures ranging from 39 to 41°C. Ocular and nasal discharges developed at day 4 with CIV89 strain and at day7 post-infection with India-Calcutta strain. Oral ulceration and necrotic lesions appeared at day 5 with CIV89 strain and at day8 with India-Calcutta. Diarrhoea was recorded in all infected goats. At day8, all infected goats were humanely slaughtered and samples were taken on autopsy for analysis.

### Uninfected goats (Control goats): No clinical signs were recorded in these control animals Clinical response of Cattle infected with isolates of PPRV from each of the four lineages

The PPRV isolates, CIV89, Nigeria 75/3, Ethiopia, India-Calcutta, representing the PPRV four lineages were used to infect young cattle (three animals / PPRV strain lineage).

Rectal temperature remained stable between 38 and 39°C during the observation period. Only one animal in the CIV89 group reached 39.7°C for 3 days. No clinical signs were recorded during the whole observation period.

### In-contact goats: No clinical signs were observed in these animals

Control Cattle: No clinical signs were recorded in these control animals as well.

### Serological response of goats to infection with PPRV isolates

The four infected goats with CIV89 and India-Calcutta respectively seroconverted at day7 and the uninfected controls remained sero negative.

### Serological response of Cattle to infection with PPRV isolates

All the infected cattle with PPRV isolates were negative from day0 to day7 post infection after analysis of the respective serum samples with the cELISA technique. At day9, 6/12 became positive, 11/12 positive at day15 and 11/12 positive at day30. One animal of group2 (cattle infected with Nigeria 75/3, lineage2) did not seroconvert. The PI values of the positive individuals in this group 2 ranged between 50 and 54% while these values were above 65% for positive animals in groups 1, 3 and 4.

The control animals remained negative (Table2).

All in-contact goats introduced in each specific box containing infected cattle with each specific lineage of PPRV remained negative.

### Detection of viral genome

All swab samples (ocular and nasal swabs) from infected cattle with PPRV and in-contact goats were analysed using the PCR technique on cDNA generated with random hexamers. This analysis found that all collected swabs were negative along with those taken from control animals (Table2).

Samples collected from slaughtered goats (infected controls) were positive by amplifying the targeted fragment of 350 bp of the NP gene.

## Discussion

PPR is a dreadful disease of sheep and goat being a real burden on the development of these species with goat being affected more severely than sheep (15). Within goat species, there is a difference in the susceptibility to PPRV between sahelian long-legged goat breed and West African dwarf goat breed from the tropical forest region with the latest more susceptible (23, 24). Conversely, PPRV is not considered as pathogenic in cattle, domestic, and wild African buffaloes (Syncerus caffer) (25) while they can seroconvert after infection with PPRV (7, 26, 27). However, high case fatality rates (96%) were reported in India in domestic buffaloes (Bubalus bubalis) and the disease was experimentally reproduced in these animals (20, 25). In Ivory-Coast, a survey on wildlife in the National game park of Comoé during the Global Rinderpest Eradication Programme (GREP) revealed that 1/56 serum samples and three pools of five swabs samples each collected from African wild buffaloes (*Syncerus caffer*) were positive to PPRV (28). This national park harbors some villages having domestic sheep and goats and contacts with wild ruminants are frequent which contribute to cross-species transmission of PPRV.

No other cases have been reported from India since then or elsewhere in Africa in cattle or African buffaloes populations. Our study was designed to give an answer to the infection of cattle with PPRV and to demonstrate whether cattle can play an epidemiological role in the spread of PPRV infection among cattle and small ruminants’ populations. Previous study implemented in Africa with PPR virus strains from each lineage demonstrated that CIV 89 (Lineage 1) strain is highly virulent followed by India-Calcutta (Lineage4 then Ethiopia (Lineage3) and finally Nigeria 75/3 strains (Lineage2) (24). In our study, control goats challenged with CIV89 and India-Calcutta strains developped clinical signs consistent with PPR and were humanely sacrified at day8 post infection, which confirmed the virulence of PPR virus strains used in this experiment. In addition, laboratory analysis on samples collected from these animals confirmed the disease. Infection of cattle with PPRV strains from each lineage did not show any clinical signs during the observation period of 30 days along with the in-contact naive goats introduced in the respective boxes like the control cattle and non-challenged control goats. This result demonstrated that cattle, after infection with PPRV, there is no replication, at least at the level of the epithelial cells (no investigation of the others cells such as PBMC) and do not excrete the virus able to contaminate animals in close contact such as goats placed in the same box. The absence of viral excretion from these challenged cattle is confirmed by the negative results of the collected swabs using the RT-PCR technique. Furthermore, recently, authors carried out an experimentally infection of calves with PPRV and could demonstrate the presence of PPRV antigen and nucleic acid in blood, plasma and PBMCs during a long period. They concluded that cattle pose no risk in transmitting the disease as virus was absent of the natural secretion of the animals (29).

Analysis of the serum samples revealed a serconversion from day9 post-infection with 6 positive cattle out of 12, in group1 with CIV89, group3 with Ethiopia and group4 with India-Calcutta strains respectively. At day15, all animals in these groups 1, 3 and 4 became positive (9/12) and at day30 post-infection, these animals remained positive. However, only 2/3 animals, challenged cattle in group2 with Nigeria 75/3 did seroconvert. The in-contact goats remained seronegative. Our study showed that, even though there is no viral excretion, the challenged animals could elicit specific anti-PPR antibodies.

These findings from the infected cattle confirmed previous studies where cattle developed specific humoral response and the production of antibodies to naturally or experimentally infection with PPRV (10, 29-33) or with the PPR vaccine (25, 34). Furthermore, these data confirm what is observed in rural communities where small ruminants and cattle co-exist, grazing together on the same pasture. In consequence, cross-species transmission of PPRV from small-ruminants to cattle is likely to occur frequently (4). At day7 post-infection, none of cattle responded serologically to the challenge with PPRV while sheep and goats seroconvert earlier, at day7 post infection or after vaccination (24). The weak seroconversion of animals in group2 with Nigeria 75/3 strain (2/3 positive animals with PI values just above the threshold) seems to be likely linked to the virulence of the strain of PPRV. Indeed, challenged animals with strains from lineages 1, 3 and 4 induced a correct production of specific antibodies against PPRV. A study revealed that challenged goats with this PPRV strain 75/3 survived after showing mild to inapparent PPR disease and seroconverted (24). The present results from group2 confirm previous study where 66 animals seroconverted out of 93 (71%) young cattle vaccinated with the PPR vaccine 75/1. A second vaccination was carried out on the 27 negative animals (93-66) to obtain 100% positive animals (34).

We have demontrated that cattle challenged with wild-type PPRV from each lineage do not excrete the virus in the environment to contaminate in-contact animals. However these animals seroconvert following a challenge with virultent wild-types PPRV. Therefore cattle cannot be considered as a PPRV reservoir and do not play an epidemiological role in the maintenance and spread of PPRV among cattle and small rumiant’s populations. Cattle are regarded as dead end host for PPRV and can rather serve as indicators of PPRV circulation and useful animal population for surveillance in the contexte of PPR eradication programme. The results of this study are of importance to be taken into account in the current PPR global eradication programme.

## Acknowledgements

We greatly thank the Centre for Research in Agronomy (CNRA-La Mé) for their support in the implementation of the experiment.

This work was funded by a Grant fro FAO / IAEA (Contract N°5380 170 3103 E3000321. Pers No: 117744.

---

**Table 1:** Infection of Cattle with each wild type candidate from the four PPRV lineages

|  | Group1-Box1 | Group2-Box2 | Group3 - Box3 | Group4 - Box4 | Group5-Box5 |
| --- | --- | --- | --- | --- | --- |
| PPRV strains | Lineage1 : | Lineage2 : | Lineage3 : | Lineage4 : | Control Cattle |
| Species | CIV89 | Nigeria 75/3 | Ethiopia | India-Calcutta |  |
| Cattle | 732 | 743 | 752 | 695 | 780 |
|  | 761 | 764 | 772 | 741 | 781 |
|  | 772 | 782 | 776 | 767 | 785 |
| 24h after infection : Introduction of naive in-contact goats |  |  |  |  |  |
| In-contact | 1.1 | 2.1 | 3.1 | 4.1 |  |
| Goats | 1.2 | 2.2 | 3.2 | 4.2 |  |
| Separated building |  |  |  |  |  |
| Infected | Box1 : |  |  | Box2 : |  |
| Control goats | CCIV1* |  |  | CInd1** |  |
|  | CCIV2 |  |  | CInd2 |  |
| Control Naïve goats | CN1***, CN2, CN3 : Uninfected control goats in box 4 |  |  |  |  |

(\*) : Control goat infected with CIV89.
(\*\*) : Control goatinfected with India Calcutta.
(\*\*\*) : Control naive goat : no challenge with any PPRV strain.

**Table 2:**
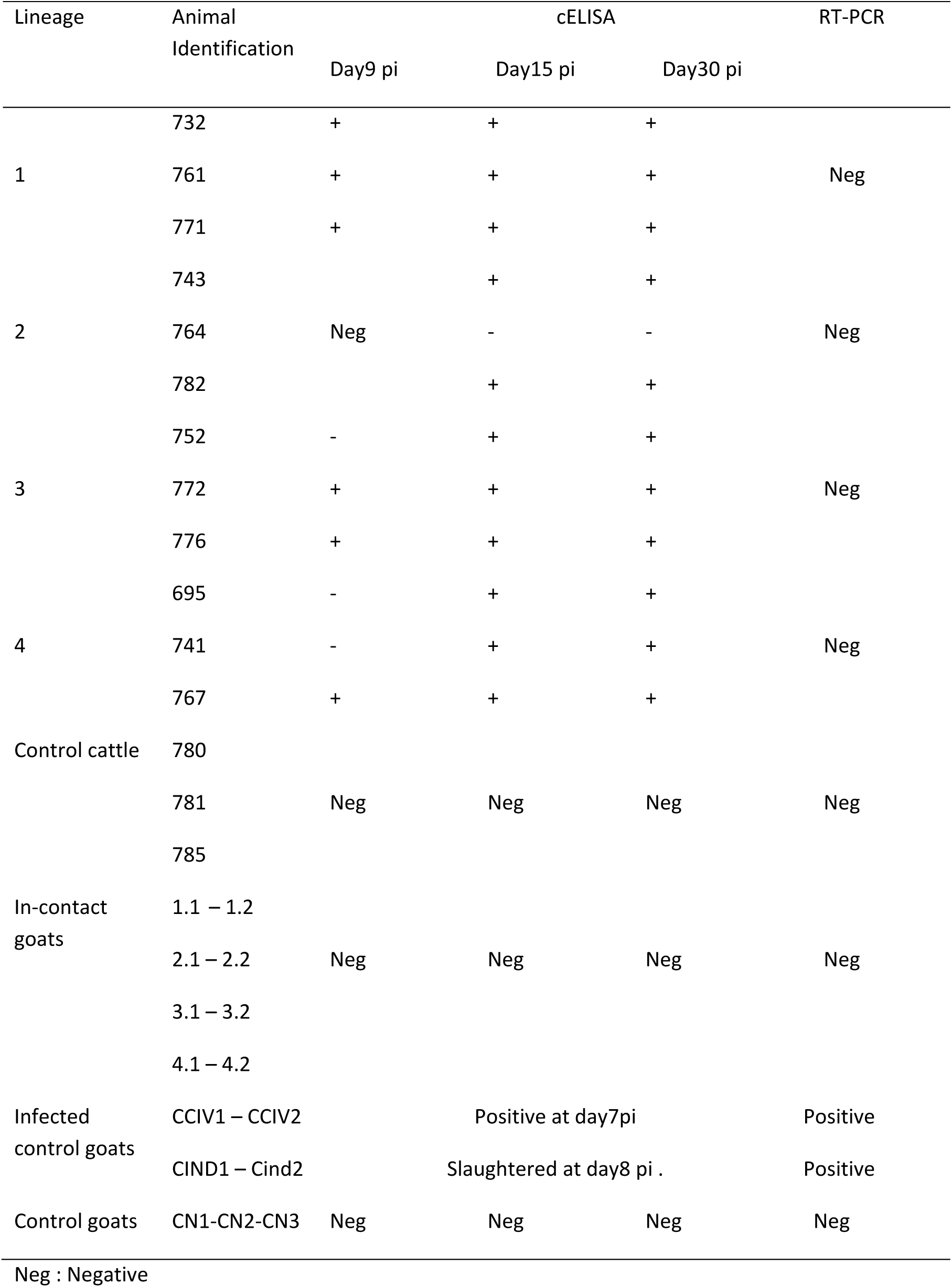
PPR specific antibodies and genome detection Results after infection of cattle with wild type of PPRV

